# Nitroxoline as a promising alternative drug for the treatment of Lyme disease based on an *in-vitro* study

**DOI:** 10.1101/2021.02.04.429852

**Authors:** Hector S. Alvarez-Manzo, Yumin Zhang, Wanliang Shi, Ying Zhang

**Affiliations:** Department of Molecular Microbiology and Immunology, Bloomberg School of Public Health, Johns Hopkins University, Baltimore, MD 21205, USA; (H.S.A.-M.); (YM.Z.); (W.S.); (H.S.A.-M.); (YM.Z.); (W.S.); State Key Laboratory for the Diagnosis and Treatment of Infectious Diseases, The First Affiliated Hospital, Zhejiang University School of Medicine, Hangzhou, Zhejiang 310003, China

**Keywords:** Lyme disease, PTLDS, Borrelia burgdorferi, nitroxoline, drug repurposing

## Abstract

Lyme disease (LD) is the most common vector-borne disease in USA and Europe and is caused by *Borrelia burgdorferi*. Despite proper treatment, approximately one fifth of patients will develop post-treatment LD syndrome (PTLDS), a condition which is poorly understood. One of the possible causes is thought to be due to persister forms of *B. burgdorferi* that are not effectively killed by the current Lyme antibiotics. In this study, we evaluated nitroxoline, an antibiotic used to treat urinary tract infections, for its activity against a stationary-phase culture enriched with persister forms of B. burgdorferi. Nitroxoline was found to be equivalent in activity against *B. burgdorferi* to cefuroxime (standard Lyme antibiotic) in different experiments. Moreover, we found that the three-drug combination cefuroxime + nitroxoline + clarithromycin eradicated 98.3% of stationary phase bacteria in the drug-exposure experiment and prevented the regrowth in the subculture study after drug exposure, as well as two-drug combinations cefuroxime + nitroxoline and clarithromycin + nitroxoline. These drug combinations should be further evaluated in a LD mouse model to assess if eradication of persister forms of B. burgdorferi *in-vivo* is possible and if so, whether nitroxoline could be repurposed as an alternative drug for the treatment of LD.

## Introduction

Lyme disease (LD) is caused by the spirochete *Borrelia burgdorferi* sensu lato complex and the spectrum of clinical manifestations is wide, ranging from mild symptoms like fatigue, headache and fever, to severe manifestations such as arthritis, carditis, encephalopathy, encephalomyelitis, and neuropathy. The number of reported cases has increased significantly in the last two decades [1]. Nowadays, there are around 30,000 cases reported in USA every year [1] and in Europe, the estimated number of cases is around 65,000 [2] with the majority in Germany [3], Austria, Lithuania, Slovenia and Sweden [4]. Nevertheless, since LD is difficult to diagnose, it is commonly underreported [5] and total numbers could be at least 10 times higher [6,7]. For this reason, it is considered a major public health problem and the most important vector-borne disease in North America [4] and Europe [2]. Treatment for the early-stage disease is administered orally with doxycycline, amoxicillin or cefuroxime. In comparison, intravenous ceftriaxone is the preferred choice for late stage manifestations [8,9]. Unfortunately, antibiotic treatment with these drugs fails in 10-20% of cases [10,11]. As a result, patients report fatigue, musculoskeletal pain [10] and some neurological symptoms like headaches and memory loss [12]. This condition is called “post-treatment Lyme disease syndrome” (PTLDS) and patients affected will see their quality of life diminished [12] with symptoms lasting for at least 6 months or longer after treatment [10,13]. PTLDS is nowadays a public health problem in USA and mathematical models calculate that the cumulative prevalence of PTLDS patients is actually around 2 million cases by year 2020 [14]. While the cause for this condition is unclear, one of the theories is that PTLDS could be due to persistent infection with persister forms of *B. burgdorferi* [10,15]. Several *in-vivo* animal studies have led to this theory by different techniques [16–18], and there is even a study that reported the presence of *B. burgdorferi* aggregates in the midgut of ticks after bloodfeeding with live-imaging, suggesting that infection with other forms than the spirochetal form *in-vivo* is possible [19].

*In-vitro* [20–23] and *in-vivo* [15] studies have shown that persister forms of *B. burgdorferi* (round-body and microcolony forms) are not easily cleared by the current Lyme antibiotics. Instead, we identified daptomycin as an antibiotic with anti-persister activity against *B. burgdorferi* and that the three-drug combination of cefuroxime + doxycycline + daptomycin is the most effective drug combination available for killing *B. burgdorferi* persisters both *in-vitro* [24,25] and *in-vivo* [15]. However, since daptomycin is an expensive parenteral drug, it is important to identify other drugs that are both more economically accessible and that can be orally administered.

In our recent study, we found that nitroxoline as a single drug, is equally effective against *B. burgdorferi* stationary-phase cultures enriched with persister forms, when compared with cefuroxime [26]. Nitroxoline (8-hydroxy-5-nitroquinoline) is an FDA-approved oral bacteriostatic antibiotic [27,28] that is commercially available in Europe [29,30] and has been used in Germany and other eastern European countries for the treatment of urinary tract infections [31]. However, in recent years, there is interest in the repurposing of this antibiotic as an anticancer drug [32–36] and also for the treatment of other infections [37–39]. Thus, our previous *in-vitro* results in addition to the studies that are looking into the possibility of repurposing this antibiotic, support the importance of performing further drug combination testing with nitroxoline in order to establish if this drug could be a useful candidate for the treatment of LD. To our knowledge, this is the first study that evaluates nitroxoline against a stationary-phase culture enriched with persister forms of *B. burgdorferi*.

## Materials and Methods

### Strain, media and culture

Barbour-Stoenner-Kelly-H (BSK-H) medium (HiMedia Laboratories Pvt Ltd.) was filter-sterilized with 0.2 μm filters and supplemented with 6% rabbit serum (Sigma-Aldrich, St. Louis, MO, USA) for the culturing of B. burgdorferi N40 strain. Cultures were incubated in 15 mL conical tubes in a microaerophilic incubator (33 °C, 5% CO2) for 7-10 days in order to reach stationary phase, equivalent to 107-108 spirochetes/mL [20]. For the evaluation of drugs, a 96-well plate was used. The stationary-phase culture (100 μL) was added to each well of the 96 well plate and drugs were added at the desired concentration. Afterwards, the plates were sealed and placed in a microaerophilic incubator for seven days at 33 °C.

### Drugs

Drugs used in this study were purchased from Sigma-Aldrich (St. Louis, MO, USA) and were dissolved in the appropriate solvents. The following drugs were utilized based on our previous studies [26]: Artemisinin (Arte), Cefuroxime (CefU), Clarithromycin (Clari), Clofazimine (CFZ), Daptomycin (Dapto), Doxycycline (Doxy), Erythromycin (Ery), Linezolid (LNZ), Nitazoxanide (NTZ), Nitroxoline (NTX), Rifabutin (Ribu). We included only drugs approved by the FDA and that can be administered orally (with the exception of Dapto which is a parenteral drug used in combination with CefU and Doxy as the three-drug control). The decision to utilize this drug is based on previous studies conducted by our research group that showed its high anti-borrelia persister activity [20,21,26]. Stock solutions were stored at −20 °C.

### Microscopy

B. burgdorferi samples were stained with the SYBR Green I/ Propidium Iodide (PI) assay and the bacterial viability was calculated with the green/red fluorescence ratio as previously described [40]. The SYBR Green I/ PI stock solution consisted of 10 μL SYBR Green I (10,000x stock, Invitrogen) and 30 μL PI (20 μM, Sigma Aldrich) into 960 μL sterile dH2O. For the work solution, the stock was diluted in a 1:4 ratio and for this study in particular, 20 μL of the SYBR Green I/ PI work solution and 40 μL of sample were added to each well to image under a BZ-X710 all-in-one fluorescence microscope (KEYENCE, Itasca, IL, USA). Three biological replicates were assessed for each drug combination and a representative picture for each sample was acquired. For the analysis of the samples, the ImageJ program was used in order to facilitate the counting of bacterial cells stained green or red and the bacterial viability of each sample was calculated by dividing the green fluorescence over the green and red fluorescence (Green/(Green + Red)).

### Drug susceptibility testing

Minimum inhibitory concentration (MIC) was determined with the microdilution method and bacterial growth inhibition was assessed after 7 days of drug addition by fluorescence microscope, after samples were stained with SYBR Green/PI. For the dose-response curves with CefU or NTX, a 7-day old *B. burgdorferi* stationary-phase culture was used and drugs were added at the desired concentration and incubated for 7 days. After this period of time, bacterial viability was assessed using the SYBR Green/PI assay. For both drugs, the Cmax concentration (Table 1) was used as the reference to choose the rest of the doses needed for the dose-response curve. Thus, in the case of CefU, the following concentrations were evaluated: untreated, 0.7, 1.75, 3.5, 7.0 (Cmax), 17.5, 35 and 70 μg/mL. For NTX, following concentrations were assessed: untreated, 0.56, 1.4, 2.8, 5.6 (Cmax), 14, 28 and 56 μg/mL.

**Table 1.**
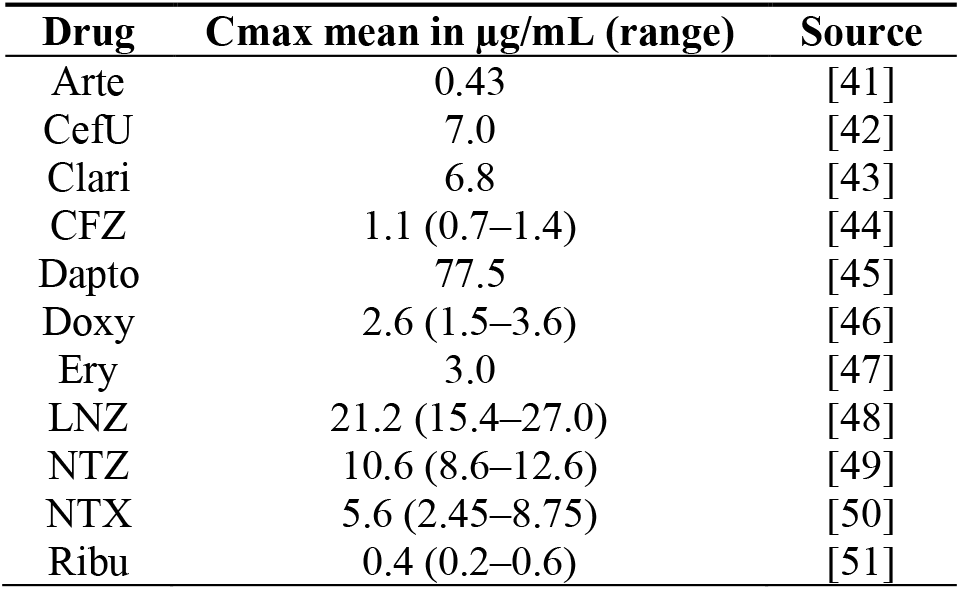
Cmax concentrations for drugs utilized in this study. In some cases, the concentration of drugs was converted from micromolar to μg/mL. In this study drugs were tested at their mean Cmax value.

| Drug | C<sub>max</sub> mean in µg/mL (range) | Source |
| --- | --- | --- |
| Arte | 0.43 | [41] |
| CefU | 7.0 | [42] |
| Clari | 6.8 | [43] |
| CFZ | 1.1 (0.7–1.4) | [44] |
| Dapto | 77.5 | [45] |
| Doxy | 2.6 (1.5–3.6) | [46] |
| Ery | 3.0 | [47] |
| LNZ | 21.2 (15.4–27.0) | [48] |
| NTZ | 10.6 (8.6–12.6) | [49] |
| NTX | 5.6 (2.45–8.75) | [50] |
| Ribu | 0.4 (0.2–0.6) | [51] |

Drug-exposure experiments with drug combinations were assessed in the same way. We used a standard concentration of 5 μg/mL as a close average Cmax concentration based on the literature, and selected drug combinations were further assessed at Cmax concentrations (Table 1). Moreover, the drug combination CefU + Doxy + Dapto was used as the positive drug control based on previous studies that showed that the combination with a cephalosporin + Doxy + Dapto is able to eradicate all borrelia bacteria including persister forms, it inhibits regrowth in the subculture study and achieves the best results in the drug-exposure experiments [23–26]. It is important to mention that Cmax concentrations were used to simulate the peak concentration of drugs in serum after oral ingestion.

### Subculture study

Selected drug combinations were also evaluated in a subculture study to prove bacterial eradication. For this part, drug combinations were added at Cmax concentration to 1 mL of a 7-day old *B. burgdorferi* culture and placed in the incubator for 7 days. Then, samples were spun down twice to remove the antibiotic. After the second wash, cell pellet was resuspended with 500 μL fresh BSK-H medium and 50 μL were taken to inoculate 1 mL fresh BSK-H to proceed with the subculture as previously described [52]. After 3 weeks, samples were evaluated for bacterial regrowth with the SYBR Green/PI assay under the microscope. All combinations tested in the subculture study were assessed in triplicates.

### Statistical analysis

The statistical analysis was carried out with the GraphPad Prism 9 software. In the case of the drug-exposure experiments, single drugs were compared with CefU as the positive drug control. For two-drug combinations, the positive drug control taken in this study was CefU + Doxy. As mentioned before, the positive three-drug control was CefU + Doxy + Dapto. The Kruskal-Wallis test was used to determine statistical differences and a p-value of less than 0.05 was considered significant.

## Results

### MIC testing and dose-response curves

The first aim of this study was to determine the MIC value for nitroxoline (NTX) and other control drugs (Table 2). The concentration that inhibited visible bacterial growth in the case of NTX was 1.25 μg/mL. In the case of cefuroxime (CefU), the MIC value was 0.08 μg/mL. We also determined the IC50 for both drugs in a 7-day old stationary-phase culture of B. burgdorferi (Figure 1). The IC50 for CefU was of 7.9 μg/mL and in comparison, NTX IC50 value was 5.3 μg/mL.

**Table 2.** MIC values of NTX and control drugs. The MIC for each drug was tested with the microdilution method.

| Drug | MIC (μg/mL) | Drug | MIC (μg/mL) |
| --- | --- | --- | --- |
| CefU | 0.08 | CFZ | 5.0 |
| Doxy | 0.15 | Dapto | >10.0 |
| NTX | 1.25 | Ery | 0.02 |
| Arte | 5.0 | LNZ | 5.0 |
| Azi | 0.6 | NTZ | 10.0 |
| Clari | 0.02 | Ribu | 5.0 |

**Figure 1.**
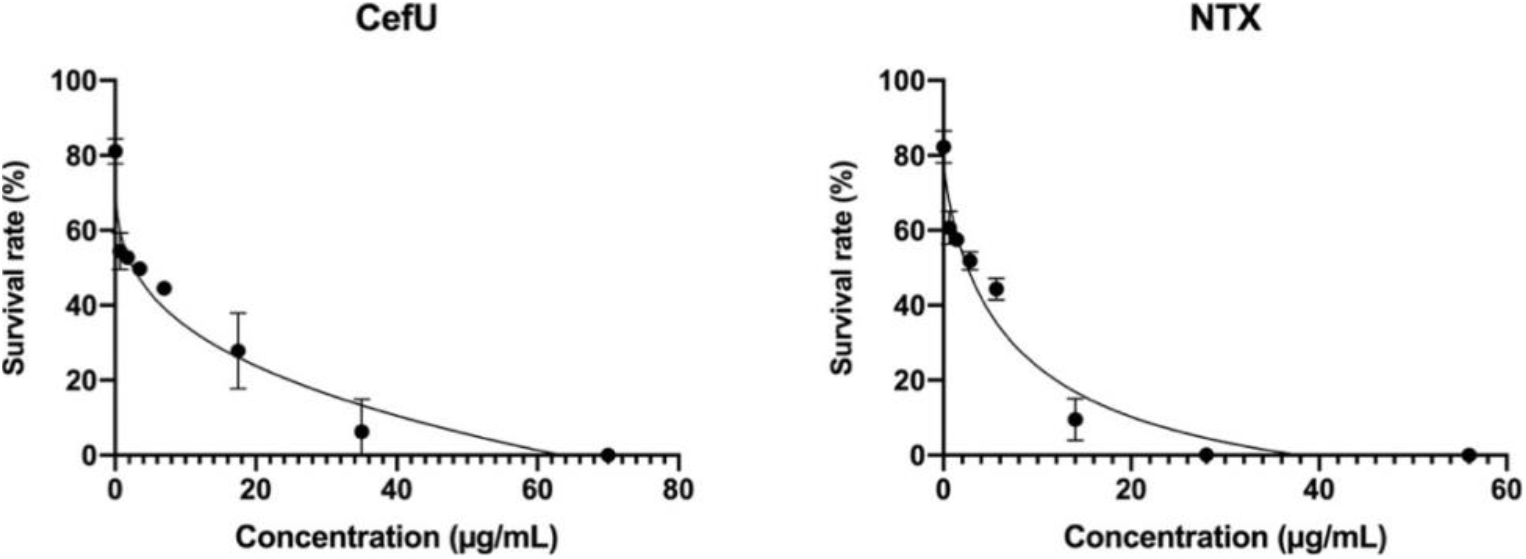
*In-vitro* drug-response curves for CefU and NTX in a 7-day old *B. burgdorferi* stationary-phase culture after a 7-day drug treatment. IC50 for CefU and NTX was 7.9 μg/mL and 5.3 μg/mL respectively. Concentrations used (μg/mL): CefU: Untreated, 0.7, 1.75, 3.5, 7.0 (Cmax), 17.5, 35, 70; NTX: Untreated, 0.56, 1.4, 2.8, 5.6 (Cmax), 14, 28, 56. Abbreviations: cefuroxime (CefU), nitroxoline (NTX).

### Activity against stationary-phase culture at 5 μg/mL and Cmax concentration

The next step was to perform a drug combination testing with nitroxoline and other drugs at a standard dose of 5 μg/mL for 7 days. Since the Cmax concentration for the different drugs is variable, we decided to take this concentration because it approaches the mean for the Cmax concentration of the drugs that we tested in this study. The stationary-phase B. burgdorferi had a survival rate of 36.3% upon 7-day treatment with Nitroxoline at 5 μg/mL. In comparison, CefU had a slightly higher survival rate of 42.3%. We then tested NTX in two- and three-drug combinations at 5 μg/mL in order to determine which combinations had a better activity against stationary-phase B. burgdorferi. We assessed a total of 19 two-drug combinations and 31 three-drug combinations at this standard concentration (Table 3). As mentioned before, the objective of this experiment was to select those concentrations that showed a promising result, in order to conduct a further experiment at Cmax concentration. Originally, we set up the threshold to determine a promising drug combination at 10% survival after treatment. Those three-drug combinations that produced bacterial viability higher than 10% would be discarded and not further evaluated. However, 87% of the combinations tested with NTX (27 out of 31) maintained bacterial viability at 10% or lower (Table 3).

**Table 3.** *B. burgdorferi* viability (in %) in a 7-day old stationary-phase culture after a 7-day drug-exposure with single drugs, two-drug and three-drug combinations with CefU or NTX at 5 μg/mL. Combinations in bold and underlined were further assessed at Cmax concentration (Table 4). A crossed line means values for these combinations were not evaluated. Abbreviations: untreated control (Cntrl), cefuroxime (CefU), doxycycline (Doxy), nitroxoline (NTX), artemisinin (Arte), azithromycin (Azi), clarithromycin (Clari), clofazimine (CFZ), erythromycin (Ery), linezolid (LNZ), nitazoxanide (NTZ), rifabutin (Ribu).

|  | Cntrl | Doxy | CefU | NTX | Arte | Azi | Clari | CFZ | Ery | LNZ | NTZ | Ribu |
| --- | --- | --- | --- | --- | --- | --- | --- | --- | --- | --- | --- | --- |
|  | 89.5 | 56.9 | 42.3 | 36.3 | 47.2 | 48.1 | 43.6 | 49.8 | 47.9 | 47.3 | 51.0 | 56.2 |
| **CefU** |  | 42.5 | — | 23.5 | 38.3 | 39.9 | 37.3 | 37.9 | 40.3 | 42.3 | 38.6 | 36.9 |
| **NTX** |  | 38.8 | 23.5 | — | 11.9 | 19.2 | 13.3 | 13.6 | 18.1 | 7.3 | 30.2 | 19.9 |
| **Arte + NTX** |  | — | 4.0 | — | — | 11.9 | 6.4 | 5.7 | 4.1 | 8.9 | 7.7 | 5.8 |
| **Clari + NTX** |  | — | **<u>5.5</u>** | — | — | — | — | 4.9 | — | **<u>7.6</u>** | **<u>9.0</u>** | 4.3 |
| **CFZ + NTX** |  | — | 8.3 | — | — | 6.0 | — | — | 2.5 | 7.5 | 11.4 | 5.0 |
| **LNZ + NTX** |  | — | **<u>2.7</u>** | — | — | 11.7 | — | — | 7.2 | — | **<u>6.1</u>** | 12.9 |
| **NTZ + NTX** |  | — | **<u>7.7</u>** | — | — | 8.7 | — | — | 10.0 | — | — | 8.6 |
| **Ribu + NTX** |  | — | 4.7 | — | — | 4.2 | — | — | 5.8 | — | — | — |

**Table 4.** Bacterial viability (in %) in a *B. burgdorferi* 7-day old stationary-phase culture after drug-exposure experiment at Cmax concentration after a 7-day treatment. A crossed line means values for these combinations were not evaluated. Combinations in bold and underlined were further assessed in a subculture study (Figure 4, Table 6). CefU + Clari + NTX did not show statistical significance when compared to the three-drug control CefU + Doxy + Dapto and NTX and Clari did not show statistical significance when compared to CefU. In comparison, CefU + NTX, CefU + Clari and Clari + NTX showed statistical significance when compared to CefU + Doxy. Abbreviations: untreated control (Cntrl), cefuroxime (CefU), doxycycline (Doxy), nitroxoline (NTX), clarithromycin (Clari), daptomycin (Dapto), linezolid (LNZ), nitazoxanide (NTZ).

|  | Cntrl | CefU | Doxy | NTX | Clari | LNZ | NTZ |
| --- | --- | --- | --- | --- | --- | --- | --- |
|  | 95.6 | 38.0 | 64.2 | 37.6 | 40.2 | 48.7 | 52.2 |
| **CefU** |  | — | 38.7 | 18.0* | 25.1* | 37.2 | 35.4 |
| **NTX** |  | 18.0 | 34.8 | — | 19.4* | 30.0 | 33.9 |
| **CefU + Dapto** |  | — | 0.0 | — | — | — | — |
| **CefU + NTX** |  | — | — | — | **<u>1.7</u>** | **<u>10.2</u>** | **<u>9.8</u>** |
| **Clari + NTX** |  | — | — | — | — | **<u>13.6</u>** | **<u>14.3</u>** |
| **LNZ + NTX** |  | — | — | — | — | — | **<u>21.0</u>** |

\* shows statistical significance.

Thus, we decided to modify the criteria to decide which combinations to test at Cmax concentration. Based on the literature, we chose to further evaluate drugs with a Cmax concentration higher than 5 μg/mL. In the case of CefU, NTX, Clari, LNZ and NTZ, their Cmax concentrations are above the standard dose chosen (5 μg/mL). Thus, we hypothesized that drug combinations with a Cmax above 5 μg/mL would result in a better result than at the standard dose. We did not further evaluate combinations with Arte, Azi, CFZ, Ery and Ribu, since Cmax concentrations for these drugs are below 5 μg/mL. We decided this rationale would determine which combinations to test at Cmax concentration.

We tested a total of 6 combinations at Cmax concentrations: CefU + Clari + NTX, CefU + LNZ + NTX, CefU + NTZ + NTX, Clari + LNZ + NTX, Clari + NTZ + NTX and LNZ + NTZ + NTX. Cmax concentrations for these drugs were taken from the literature and survival rates for these combinations at Cmax concentrations are depicted in Table 4. We compared the NTX three-drug combinations with the three-drug positive control CefU + Doxy + Dapto. The best combination was CefU + Clari + NTX, with a mean survival rate of 1.7% (Figure 2 and Figure 3). When performing the statistical analysis, NTX (survival rate 37.6%) and Clari (survival rate 40.2%) were not statistically significant when compared to CefU (survival rate 38.0%). CefU + NTX (survival rate 18.0%), CefU + Clari (survival rate 25.1%) and Clari + NTX (survival rate 19.4%) were statistically significant when compared to the two-drug control CefU + Doxy (survival rate 38.7%). At last, CefU + Clari + NTX (survival rate 1.7%) was not statistically significant when compared to three-drug control CefU + Doxy + Dapto (survival rate 0.0%).

**Figure 2.**
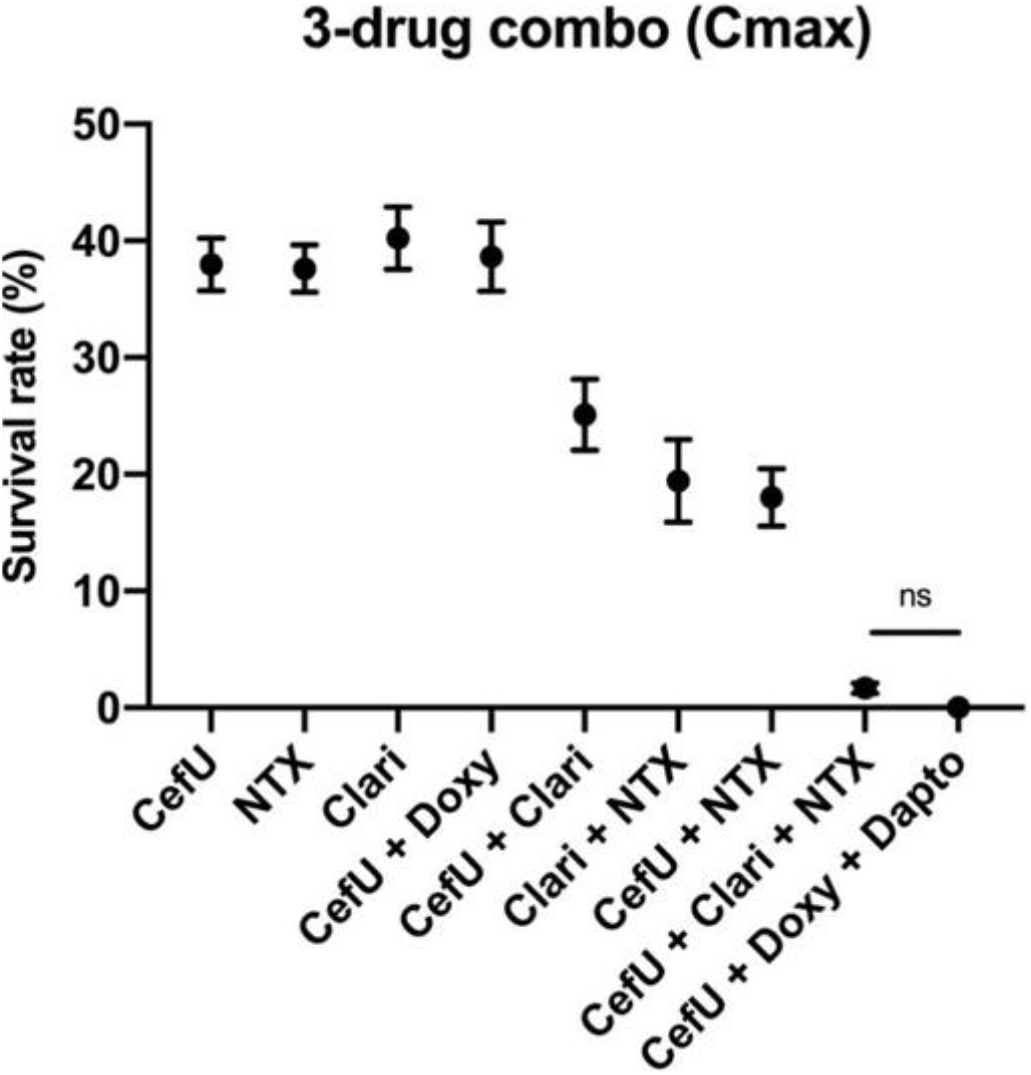
NTX-drug combinations of interest evaluated at Cmax concentration in a *B. burgdorferi* 7-day old stationary-phase culture after a 7-day treatment. CefU + Clari + NTX (survival rate 1.7%) was not statistically significant when compared to the three-drug control CefU + Doxy + Dapto (survival rate 0.0%). Abbreviations: cefuroxime (CefU), doxycycline (Doxy), nitroxoline (NTX), clarithromycin (Clari), daptomycin (Dapto). ^ns^ no statistical significance.

**Figure 3.**
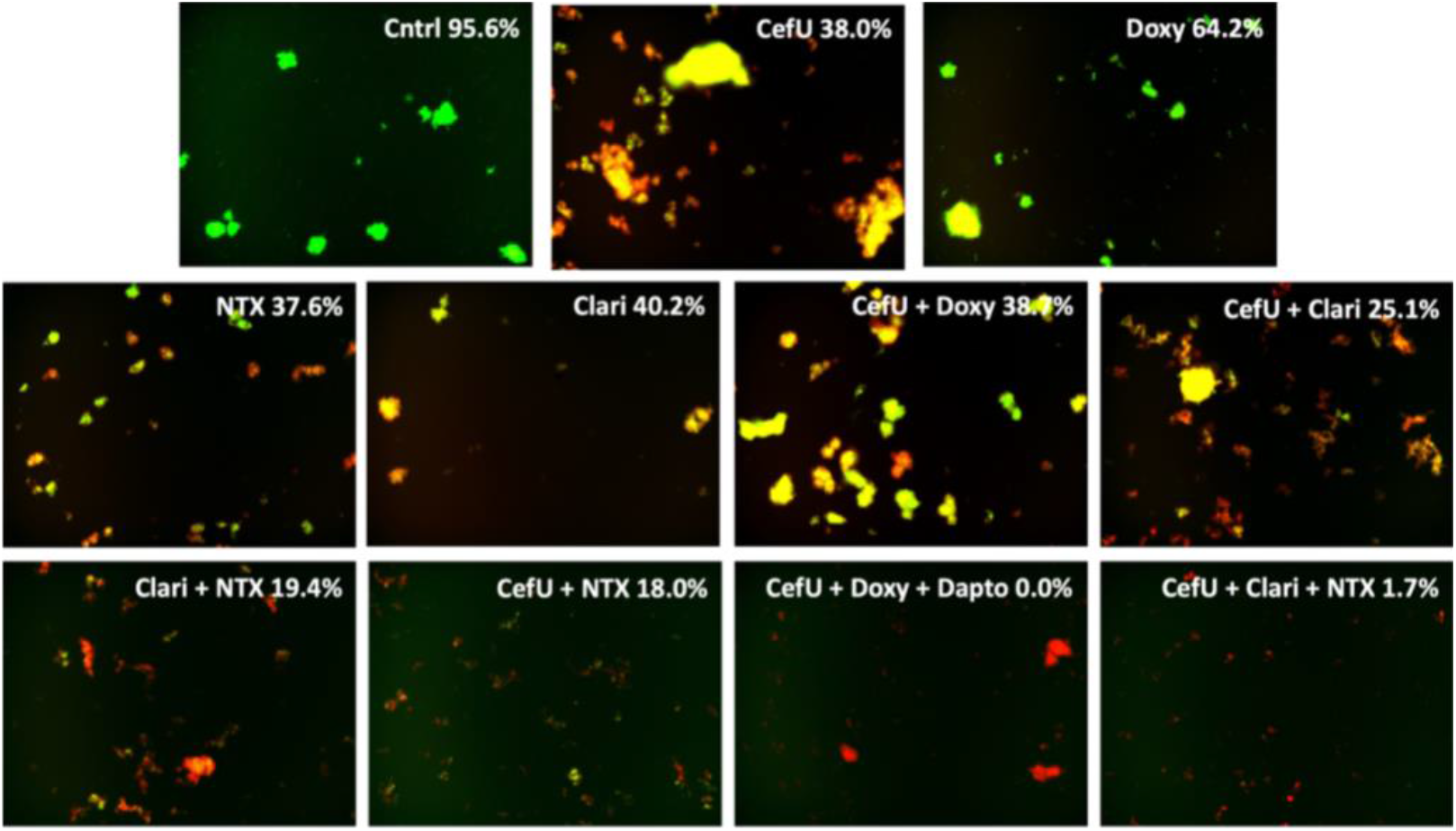
Microscope images for single drugs, two-drug and three-drug combinations of interest at Cmax concentration in a *B. burgdorferi* 7-day old stationary-phase culture after a 7-day treatment. Abbreviations: untreated control (Cntrl), cefuroxime (CefU), doxycycline (Doxy), nitroxoline (NTX), clarithromycin (Clari), daptomycin (Dapto).

**Figure 4.**
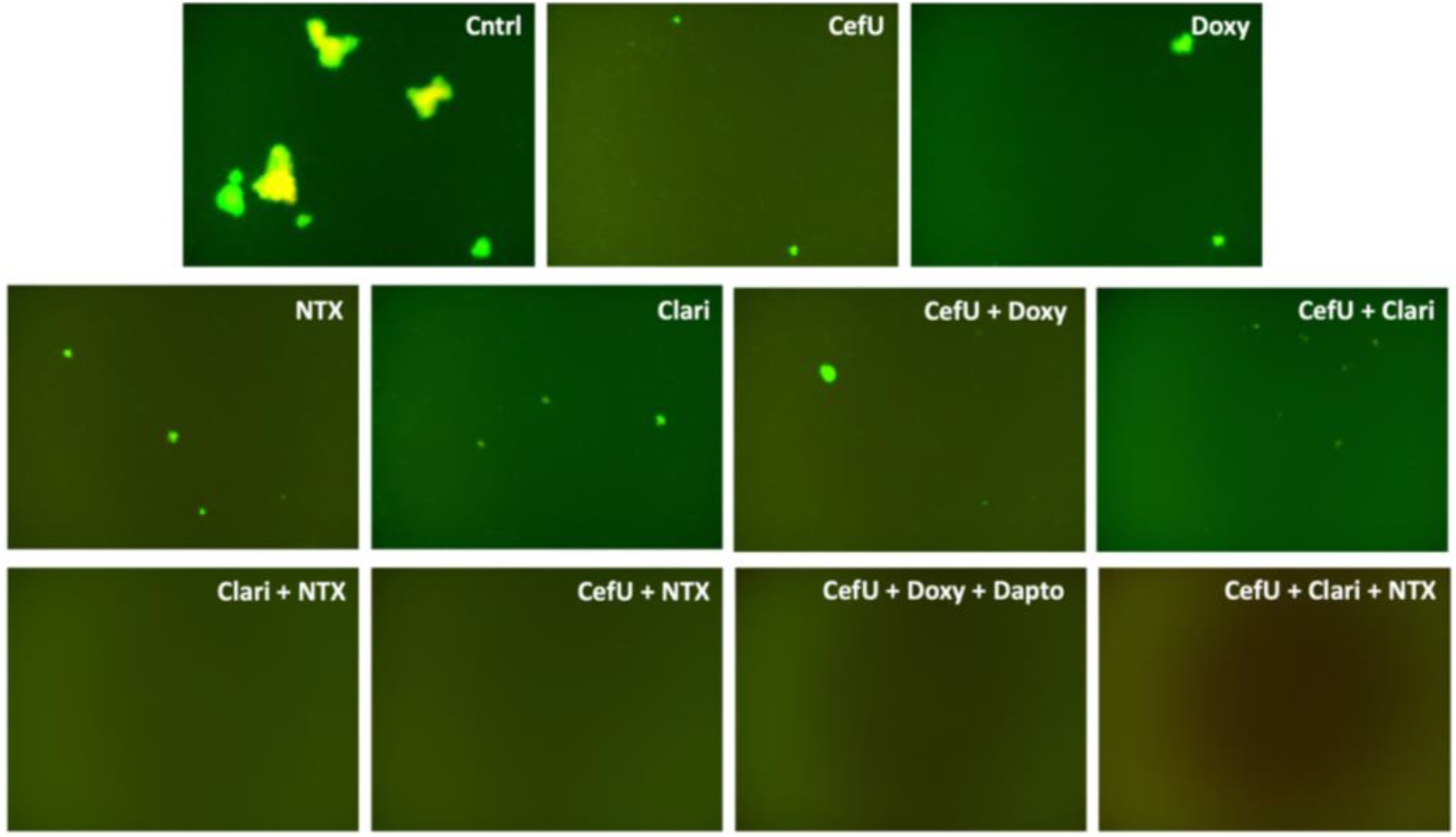
Microscope images for subcultures of single drugs, two-drug and three-drug combinations of interest. Abbreviations: untreated control (Cntrl), cefuroxime (CefU), doxycycline (Doxy), nitroxoline (NTX), clarithromycin (Clari), daptomycin (Dapto).

We then sought to compare the effect of drug combinations with NTX in comparison with drug combinations with CefU (standard Lyme antibiotic used for early disseminated and late disseminated disease) at Cmax concentrations. Thus, we chose those combinations tested at Cmax concentration that did not include CefU (Clari + LNZ + NTX, Clari + NTZ + NTX and LNZ + NTZ + NTX), replaced NTX with CefU and tested the following combinations: Clari + LNZ + CefU, Clari + NTZ + CefU and LNZ + NTZ + CefU. Survival rates for the comparison of these three-drug combinations are in Table 5. All six combinations, either with CefU or with NTX, showed to be better combinations when compared to the respective control (Table 5, left column). The best drug combinations were Clari + NTZ + CefU, Clari + LNZ + NTX and Clari + NTZ + NTX, since these kept bacterial viability at 12.6%, 13.6% and 14.3% respectively.

**Table 5.** Drug-exposure experiment with single drugs, two-drug and three-drug combinations directly comparing bacterial viability (in %) of CefU and NTX in a 7-day old *B. burgdorferi* stationary-phase culture. A statistical analysis was performed comparing the performance of combinations with CefU and NTX to the control column. All combinations with CefU and NTX were statistically significant when compared to the respective control (left column). Abbreviations: untreated control (Cntrl), cefuroxime (CefU), nitroxoline (NTX), clarithromycin (Clari), linezolid (LNZ), nitazoxanide (NTZ).

|  | Cntrl | CefU | NTX |
| --- | --- | --- | --- |
|  | 95.6 | 38.0* | 37.6* |
| **Clari** | 40.2 | 25.1* | 19.4* |
| **LNZ** | 48.7 | 37.2* | 30.0* |
| **NTZ** | 52.2 | 35.4* | 33.9* |
| **Clari + LNZ** | 36.9 | 17.7* | 13.6* |
| **Clari + NTZ** | 42.0 | 12.6* | 14.3* |
| **LNZ + NTZ** | 46.8 | 38.1* | 21.0* |

\* shows statistical significance.

### Subculture study

At last, we performed a subculture study to ensure bacterial killing by drug combinations (Figure 4, Table 6). We only tested CefU + Clari + NTX, since it was the only combination that achieved the best activity among other drug combinations tested with no statistical difference when compared to the three-drug positive control CefU + Doxy + Dapto. Both three-drug combinations achieved total eradication in the subculture study. In addition, the two drug combinations CefU + NTX and Clari + NTX also achieved eradication in this assay, despite not showing a very low survival ratio in the drug-exposure experiment (18.0% and 19.4% respectively). However, single drugs Doxy, Cefu, Clari, NTX, and the two-drug controls CefU + Doxy and CefU + Clari showed regrowth after 3 weeks subculture study (Figure 4, Table 6).

**Table 6.** Results of a 3-week subculture study for a 7-day old B. burgdorferi stationary-phase culture following a 7-day drug treatment at Cmax concentrations. Subcultures were assessed in triplicates. Each “+” (regrowth) or “–” (no regrowth) symbol represents the result for one replicate. CefU + NTX, Clari + NTX, CefU + Doxy + Dapto and CefU + Clari + NTX did not regrow after 3 weeks in the subculture experiment. Abbreviations: untreated control (Cntrl), cefuroxime (CefU), doxycycline (Doxy), nitroxoline (NTX), clarithromycin (Clari), daptomycin (Dapto), linezolid (LNZ), nitazoxanide (NTZ).

| Drug | Subculture results | Drug | Subculture results |
| --- | --- | --- | --- |
| Cntrl | +++ | CefU + NTX | --- |
| CefU | +++ | Clari + NTX | --- |
| Doxy | +++ | CefU + Clari + NTX | --- |
| NTX | +++ | CefU + Doxy + Dapto | --- |
| Clari | +++ |  |  |
| CefU + Doxy | +++ |  |  |
| CefU + Clari | +++ |  |  |

## Discussion

We evaluated the performance of NTX compared to CefU at different concentrations against *B. burgdorferi* stationary-phase cultures. The goal of our study was to determine the *in-vitro* performance of NTX and compare it to the performance of the standard Lyme antibiotic CefU. The MIC test showed the MIC value for NTX was 1.25 μg/mL, consistent with our previous finding [26], while CefU and Doxy MIC-values were 0.08 μg/mL for both, also consistent with previous results [53]. Moreover, the NTX MIC value is below its Cmax concentration (5.6 μg/mL), indicating it could reach growth inhibitory concentration when used *in-vivo*.

We were also interested in determining the effect of NTX in *B. burgdorferi* stationary-phase cultures enriched with persisters. In a previous study conducted by our research group [26], we established that NTX at Cmax achieved a similar eradication rate of a *B. burgdorferi* stationary-phase culture when compared to CefU also at Cmax. In the present study, CefU (survival rate 38.0%) and NTX (survival rate 37.6) single-drug results in the drug-exposure experiment were consistent with our previous study [26]. Two-drug combinations CefU + NTX (survival rate 18.0%), CefU + Clari (survival rate 25.1%) and Clari + NTX (survival rate 19.4%) were the best two-drug combinations and proved to be statistically significant when compared to the two-drug control CefU + Doxy (survival rate 38.7%). In the case of the three-drug combinations assessed at Cmax (Table 4), CefU + Clari + NTX achieved a survival rate of 1.7% and showed no statistical significance compared with the three-drug control CefU + Doxy + Dapto (survival rate 0.0%). This is particularly interesting since this is the first oral combination that we have identified to be equivalent towards the three-drug combination with CefU + Doxy + Dapto. Besides, it also makes sense that this combination in particular achieved the best result, since two-drug combinations that contain this very same drugs (CefU + NTX, CefU + Clari and Clari + NTX), achieved in this study the best two-drug results as previously mentioned. Thus, the combination of these three-drugs proved to be the strongest combination tested in this study and proved to be equivalent to the positive control. Moreover, CefU + Clari + NTX was also the only combination that reduced the survival rate when adjusting the concentration to Cmax. All other three-drug combination survival rates increased when compared with their 5 μg/mL result (Table 3).

Finally, we replaced CefU with NTX in some two-drug and three-drug combinations to compare the effect of each drug and determine which one could have a higher impact on persister forms of *B. burgdorferi* (Table 5). We found that combinations with either CefU or NTX were equally effective when compared to their control. Thus, although not better, it appears at least *in-vitro* that NTX has similar activity as the standard Lyme antibiotic CefU when used alone and in drug-combinations against stationary-phase cultures enriched with persister forms of *B. burgdorferi*.

In the subculture study (Table 6), the positive control CefU + Doxy + Dapto and the combination of interest CefU + Clari + NTX were shown to completely eradicate *B. burgdorferi* stationary-phase culture as demonstrated by lack of regrowth after 3 weeks of subculturing. Interestingly, not only these combinations achieved eradication, but CefU + NTX and Clari + NTX also did. This finding in particular suggests that it is not necessary for a drug-combination in vitro to achieve complete killing in the drug-exposure experiment, to be able to achieve eradication in the subculture study.

NTX has been described as a metal-chelator drug. Specifically, an *in-vitro* study showed that NTX dispersed *Pseudomonas aeruginosa* biofilm by chelating Fe^2+^ and Zn^2+^ ions [37]. Another study found that NTX had a high antimicrobial activity against *Escherichia coli*, *Providencia rettgeri* and *Klebsiella pneumoniae* [38]. Finally, biofilms formed by methicillin-resistant *Staphylococcus aureus* and *Acinetobacter baumannii* are also eradicated by NTX *in-vitro* [39]. In the stationary-phase cultures, as seen in Figure 3 (Cntrl survival rate 95.6%), *B. burgdorferi* tends to form microcolonies, which is consistent with a previous study [24]. These *B. burgdorferi* microcolonies could be analogous to small biofilms, as *B. burgdorferi* is able to generate biofilms *in-vitro* [54]. Thus, since NTX does have a high activity against biofilms as mentioned previously, the activity of this drug against *B. burgdorferi* stationary-phase cultures is not surprising. However, it still remains to be determined as to how this drug might be eradicating *B. burgdorferi. B. burgdorferi* does not utilize iron for its metabolism. Instead, different studies have demonstrated that manganese and copper might have an important role in the metabolism of *B. burgdorferi*, since these are the cofactors necessary for molecules related to virulence [55], mammalian adaptation [56], survival [57,58], and persistence within the tick [59]. As mentioned before, NTX has shown to chelate Zn^2+^ from *P. aeruginosa* biofilm, achieving biofilm disruption *in-vitro* [37]. Thus, it is possible that NTX might be chelating zinc or other ions like manganese or copper and as a result, causing *B. burgdorferi* to die. This hypothesis still has to be proven with the addition and removal of zinc, manganese, copper and other metals from the BSK-H medium and the treatment with NTX in future studies.

NTX is an FDA-approved drug not commercially available in USA but it is clinically used in some European countries [29–31]. Cephalosporin and tetracycline allergy prevalence among the general population ranges from 0.8-1.7% and 0.5-1.6% respectively and it is more frequent among women and the elderly population [60,61]. If further investigated, NTX could be repurposed for the treatment of acute stages of LD for those patients that do not tolerate treatment with the common Lyme antibiotics. In addition, our results regarding the activity of NTX as single drug and in combination (CefU + Clari + NTX), could potentially be useful in treating PTLDS due to persister forms of *B. burgdorferi*. We realize that the main limitation of this study is that we only carried out *in-vitro* experiments. However, as far as we are concerned, this is the first study to evaluate NTX in detail against *B. burgdorferi*, and our findings are completely new, promising and warrant future studies with NTX in cell lines and in animal models of *B. burgdorferi* infection. Moreover, we only tested NTX in the context of *B. burgdorferi* sensu stricto N40 strain. It will be relevant to evaluate if NTX has activity against *Borrelia afzelii* and *Borrelia garinii*, since these are the most prevalent genospecies that cause LD in Europe [62–64], where NTX is commercially available.

## Conclusions

We showed that NTX is as active as CefU in the MIC experiment, as well as against non-growing persister forms of B. burgdorferi. We found that the three-drug combination CefU + Clari + NTX was equivalent to the previous three-drug control with CefU + Doxy + Dapto. Since the combination with NTX found in this study involves only oral drugs (unlike the combination with the intravenous drug Dapto), we propose that this new combination could be considered as the new positive control for further in-vitro studies that evaluate drugs in the context of B. burgdorferi stationary-phase cultures. Moreover, CefU + Clari + NTX, as well as two-drug combinations CefU + NTX and Clari + NTX should be investigated in the future in mouse models to establish if these combinations are able to eradicate persister forms of B. burgdorferi in-vivo. Our results suggest that NTX, if further investigated and confirmed, could be repurposed for the treatment of active stages of LD. Moreover, it could also be useful for patients with PTLDS if the origin of PTLDS is caused by the presence of persister forms of B. burgdorferi. However, studies in cell lines and in-vivo experiments have to be carried out before repurposing this drug as an alternative. Finally, the elucidation of the mechanism by which NTX eradicates B. burgdorferi is key to better understanding the survival strategies of B. burgdorferi persisters and also for identification of new antimicrobial targets.

## Author Contributions

Conceptualization, H.S.A.-M. and Y.Z. (Ying Zhang); methodology, H.S.A.-M. and Y.Z. (Yumin Zhang); formal analysis, H.S.A.-M.; data curation, H.S.A.-M.; writing—original draft preparation, H.S.A.-M.; writing—review and editing, Y.Z. (Ying Zhang); supervision, W.S.; funding acquisition, Y.Z. (Ying Zhang). All authors have read and agreed to the published version of the manuscript.

## Acknowledgments

We acknowledge the support given by the Global Lyme Alliance, the Steven and Alexandra Cohen Foundation, the LivLyme Foundation, NatCapLyme, and the Einstein-Sim Family Charitable Fund.

